# Circulating *Chlamydia trachomatis*-specific T Cell Immunity Reflects Widespread Exposure in South African Adolescents and Young Women

**DOI:** 10.1101/2025.07.10.664075

**Authors:** Rubina Bunjun, Micaela Lurie, Smritee Dabee, Shaun Barnabas, Vanessa Maseko, Shameem Z. Jaumdally, Hoyam Gamieldien, David A. Lewis, Heather B. Jaspan, Katherine Gill, Linda-Gail Bekker, Jo-Ann S. Passmore

## Abstract

*Chlamydia trachomatis* remains the most prevalent bacterial sexually transmitted infection worldwide, with adolescent girls and young women (AGYW) disproportionately affected. Despite this, vaccine development is hindered by our limited understanding of protective immunity, particularly in the context of recurrent infections and immune-mediated pathology. Here, we characterised mucosal inflammation and systemic immune responses to *C. trachomatis* in South African AGYW (n=145), stratified by exposure history based on nucleic acid amplification testing (NAAT) and anti-*C. trachomatis* IgG levels. Cervicovaginal cytokines were quantified from menstrual cup secretions, and cervical and peripheral blood T cell activation assessed by flow cytometry. CD4+ T cell responses to recombinant MOMP were measured in a subset of 46 women who were NAAT-positive and/or seropositive. Untreated or recurrent *C. trachomatis* infection (NAAT+/IgG+) was associated with increased cervical T cell activation. All women with untreated or recurrent infections had detectable circulating *C. trachomatis*-specific CD4+ T cells in blood; however, response magnitude was 2.4-fold lower than in women with cleared or primary infections. Women with cleared infection had the highest proportions of *C. trachomatis*-specific multifunctional CD4+ T cells, while those with untreated or recurrent infections had almost none. Notably, systemic *C. trachomatis*-specific Th1 responses were inversely correlated with genital tract concentrations of inflammatory cytokines including IL-1β, TNF, and IL-17. These findings demonstrate that both the magnitude and quality of the systemic CD4+ T cell responses are critical components of protective immunity to *C. trachomatis*, and may limit mucosal immunopathology, which has important implications for vaccine strategies and evaluation in high-risk populations.

## INTRODUCTION

*Chlamydia trachomatis* is the most prevalent bacterial sexually transmitted infection (STI) globally. In 2020, the World Health Organization (WHO) estimated that there were more than 120 million new infections worldwide [1]. Adolescent girls and young women (AGYW), in particular, experience high chlamydia burden [2]. Although the interruption in sexual healthcare delivery caused by the COVID-19 pandemic contributed to an increase in infections [3], chlamydia prevalence was already increasing in 2019 [4,5]. Despite the availability of effective antibiotic treatment, many infections remain unnoticed and untreated [6], particularly in low-and-middle income countries (LMICs), which rely on the syndromic management for STIs [7–9]. The WHO has set a target of a 50% reduction in new cases of chlamydia by 2030 [1]. **Achieving this objective requires a prevention-** oriented strategy, with an emphasis on vaccines [10].

CTH522:CAF01 is the first chlamydia vaccine to enter human trials in more than a decade, and has just completed Phase I trials [11]. Chlamydia vaccine development has been hindered by several immune-related obstacles [12,13]. Live-attenuated vaccines carry the risk of inducing pathology [14]. Immunopathology in the female genital tract is associated with pelvic inflammatory disease and subsequent infertility [15,16], and may increase the risk of HIV acquisition [17]. Therefore, an effective vaccine needs to induce a protective immune response without the accompanying mucosal immunopathology. Furthermore, determining correlates of protection is complicated by the natural history of *C. trachomatis* infection. Untreated or recurrent infections are common in young women but decrease with age, suggesting some immune protection induced by infection, although incomplete [18–21].

CD4+ T cells play a key in immunity against chlamydia, primarily due to their production of Th1 cytokines [22–24]. Clinical studies have focused on antigen mapping, with limited data characterising the T cell response to *C. trachomatis*, especially in high-risk young women or in relation to exposure history [25–28]. Consequently, there is an urgent need to better understand the development of immunity, while also considering the development of immune-mediated pathology due to multiple exposures. We characterised genital tract inflammation and systemic CD4+ T cell responses to *C. trachomatis* in South African AGYW, stratified by their infection history and relative exposure levels, to better understand immune correlates of prior cleared infections, and the effect of continuous pathogen exposure.

## RESULTS

### Cohort description

We used baseline samples from a well-characterised cohort of HIV-uninfected AGYW (median age of 18 years old) from South Africa [29–31]. Participants were recruited within three years of sexual debut and reported a median of two sexual partners (Table 1). Hormonal contraceptive use was high (n=143/145), but only 34% (42/122) of participants reported regular condom use within the past year. Importantly, high burdens of chlamydia (43%) were found, with low rates of previous STI diagnosis or treatment (14%), highlighting the need for better sexual health interventions.

**Table 1.** Participant characteristics.

| Characteristic | N=145 |
| --- | --- |
| Age in years [median (IQR)] | 18 (17-20) |
| Age of sexual debut (years) [median (IQR)] | 16 (15-17) |
| Lifetime sexual partners [median (range)] | 2 (1-13) |
| Multiple sexual partners in the last 12 months [n/N(%)] | 33/111 (30%) |
| **Contraceptive use [n/N (%)]** |  |
| Ever pregnant | 30/122 (25%) |
| Previous contraceptive use | 96/122 (79%) |
| Condom use during last sex act | 87/121 (72%) |
| Regular condom use in the last 12 months | 42/122 (34%) |
| **Hormonal contraceptives** |  |
| *DMPA-IM* | 27/145 (19%) |
| *Net-En* | 99/145 (68%) |
| *Combined oral contraceptive* | 7/145 (5%) |
| *LNG implant* | 9/145 (6%) |
| *Nuvaring* | 1/145 (0.7%) |
| **Sexually transmitted infections (STIs) [n/N (%)]** |  |
| *Chlamydia trachomatis* | 62/145 (43%) |
| *Neisseria gonorrhoeae* | 17/145 (12%) |
| *Mycoplasma genitalium* | 6/145 (4%) |
| *Trichomonas vaginalis* | 11/145 (8%) |
| Bacterial vaginosis (Nugent score 7-10) | 70/145 (48%) |
| Previous STI (diagnosed or treated) | 17/122 (14%) |

### Classification of *C. trachomatis* Infection History and Exposure Levels using NAAT and Serology

We used anti-*C. trachomatis* IgG serology to infer infection history. Thirty percent (30%; 44/145) of participants were seropositive, 14% (20/145), were borderline positive and 56% (81/145) were seronegative (**Supplementary Table S1**). Of those with a negative nucleic acid amplification test (NAAT), 72% (60/83) were also seronegative (**Figure 1A**). Similarly, 66% (41/62) of NAAT+ AGYW were also seropositive. Notably, 34% (21/62) NAAT+ AGYW were seronegative, signifying a primary infection. Women who were NAAT+ also had nearly 3-fold higher antibody levels than women who were NAAT-(**Figure 1B;** p<0.0001; medians: 20.35RU and 7.14RU, respectively), suggesting a degree of concordance between serology and NAAT. This suggests that AGYW with an active chlamydia infection were likely experiencing a recurring, persistent, or untreated infection. Given the low numbers of participants with previously treated STIs, it is most likely that AGYW with an active infection were undiagnosed and untreated, rather than having treatment failure (persistence) or repeated infections (recurring).

**Figure 1.**
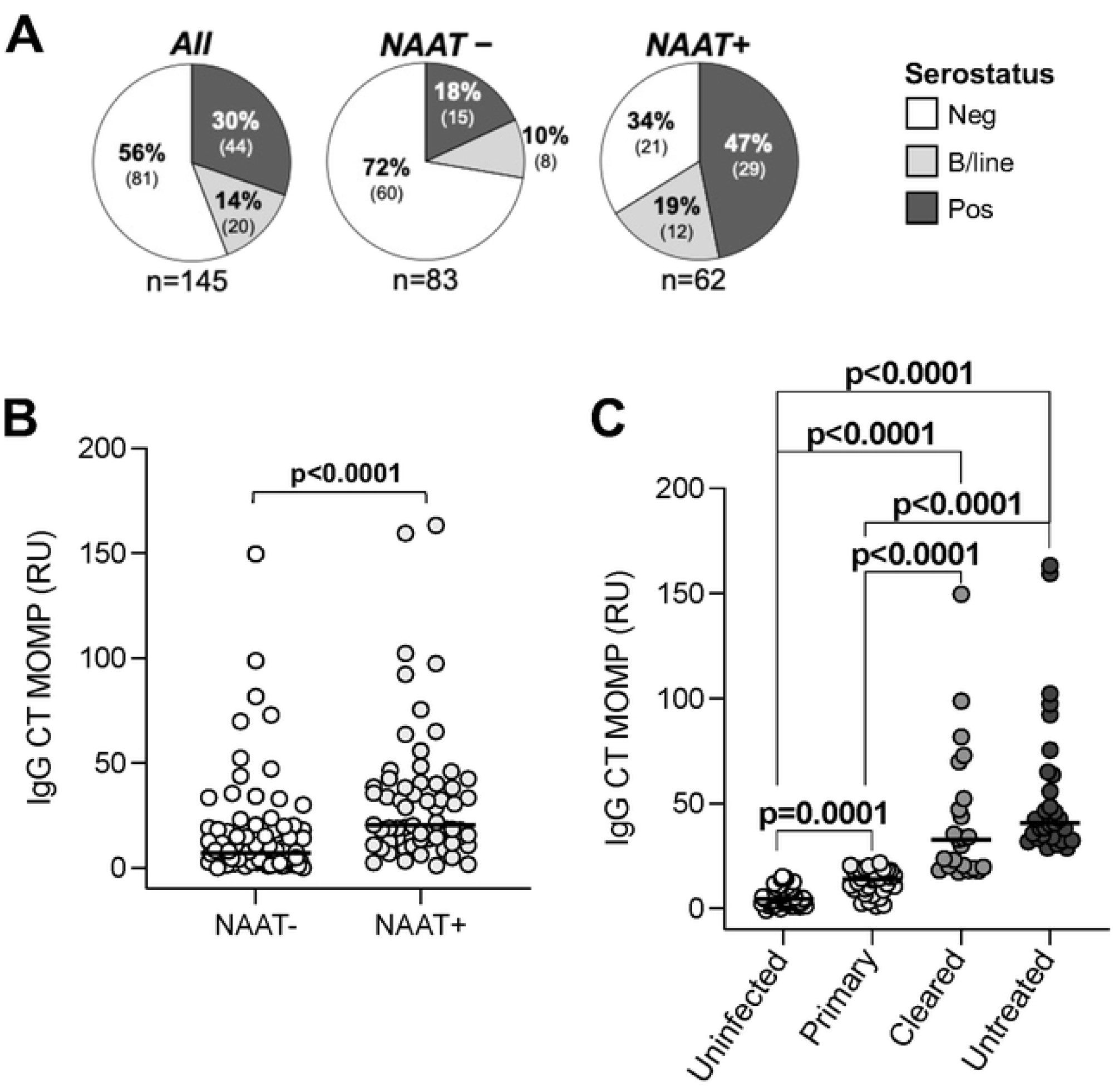
Repeat or persistent *C. trachomatis* infections were common in adolescent girls and young women (n=145) (A) Plasma serology results from the cohort measured using semi-quantitative ELISA (Euroimmun; Germany). Relative units (RU) were used to classify participants into negative (<16 RU; neg; white), borderline positive (16-21 RU; b/line; light grey) and positive (≥22 RU; pos; dark grey). (B) Distribution of IgG levels measured by ELISA. (C) IgG levels stratified by exposure. Exposure groups were assigned based on NAAT results and serostatus. Each symbol represents an individual. Data were analysed using a non-parametric Mann-Whitney U test. p<0.05 was considered statistically significant. NAAT: nucleic acid amplification test. MOMP: Major outer membrane protein. CT: *C. trachomatis*.

We stratified participants by their level of *C. trachomatis* exposure, based on NAAT and serology at baseline (**Table S1**). Participants who were negative for both were classified as uninfected, having no evidence of prior or current infection. A positive NAAT with seronegativity/borderline was indicative of primary exposure reflecting recent infection prior to detectable antibody response. AGYW with a negative NAAT who were seropositive/borderline had a cleared past infection, with clearance of the organism but persistence of antibodies. Participants who were positive for both NAAT and serology were considered to have ongoing or repeated exposure, consistent with untreated, persistent, or recurrent infection. Antibody levels increased progressively across exposure groups, with median IgG values of 4.61RU, 13.85RU, 32.98RU, and 40.69RU, respectively (**Figure 1C**). This grouping framework enabled a nuanced assessment of exposure history in a highly affected cohort and provided a unique window into the development of immune responses during early *C. trachomatis* infection.

### *C. trachomatis* Infection History and Genital Tract T Cell Activation

Multiple exposures to *C. trachomatis* have been implicated in the development of reproductive tract immunopathology [15]. Previous work within this cohort showed that *C. trachomatis* infection was associated with elevated concentrations of genital tract inflammatory cytokines including IL-6, TNF, IL-1β, and IFN-γ [32]. Building on these findings, we examined the relationships between exposure history, as measured by serology, and the immune environment in the female genital tract. Anti-chlamydia IgG positively correlated with the concentration of cervicovaginal IFN-γ (**p=0.037**, *r*= 0.174; **Figure 2A**), suggesting a significant but weak link between multiple or long-term exposure and mucosal Th1-type cytokine production. No associations were observed between antibody levels and other cytokines (IL-6, TNF, IL-1β). Notably, systemic anti-chlamydia IgG also positively correlated with the frequency of activated cervical CD4+ T cells expressing HLA-DR (p=0.018, *r*=0.211; **Figure 2B**), and those co-expressing CD38 and HLA-DR (p=0.012, *r*=0.223; Figure 2C). A similar association was seen with activated CD8+HLA-DR+ T cells (p=0.019, *r*=0.209; data not shown), though no correlations were found with CD4+CD38+ T cells or CD8+CD38+ T cell subsets alone.

**Figure 2.**
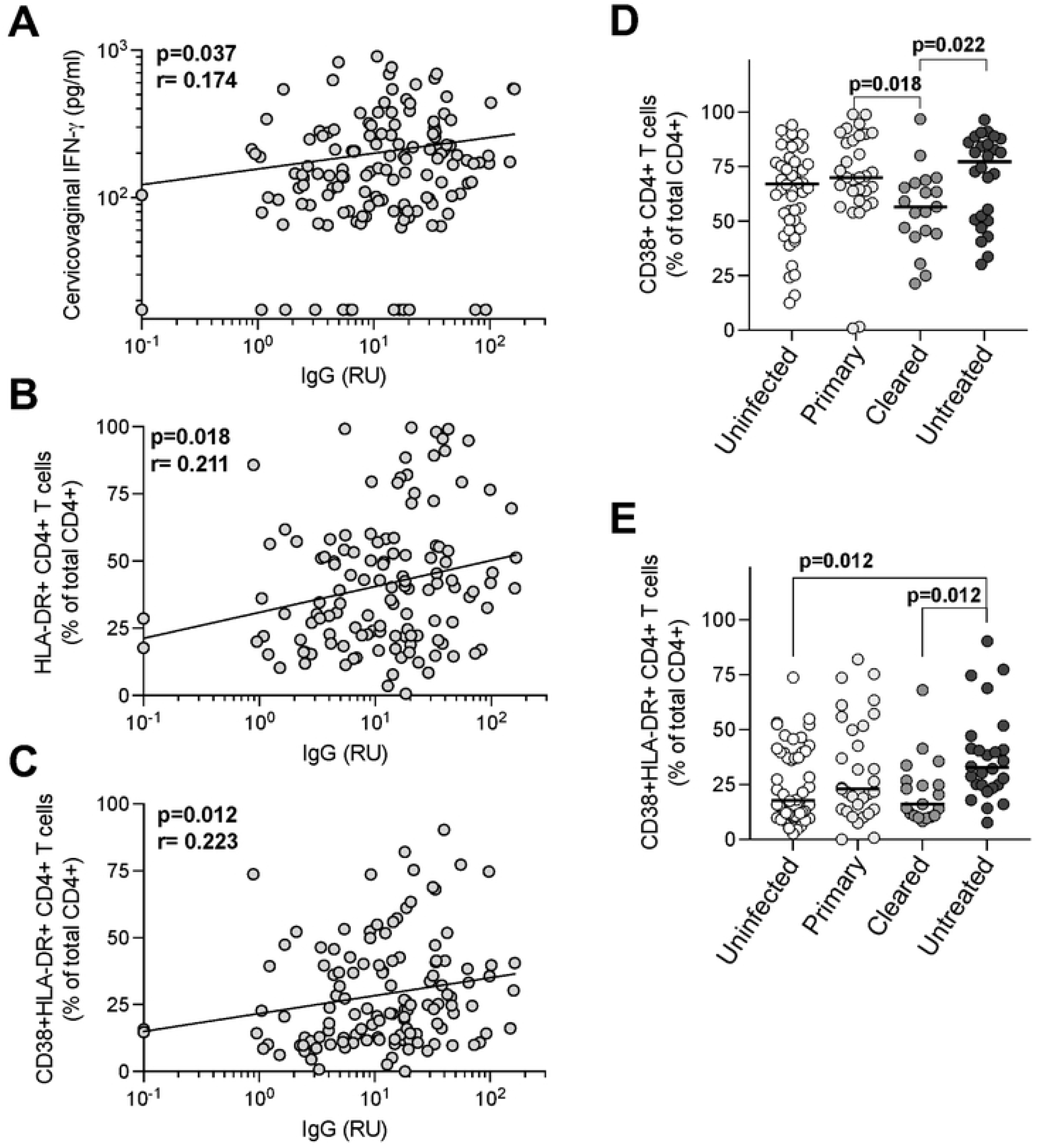
Exposure to *C. trachomatis* is associated with T cell activation in the female genital tract. (A) Associations between IgG and Cervicovaginal IFN-γ concentrations (n=144) and (B-C) frequencies of activated cervical T cells (n=126). (D-E) Associations between exposure score and the frequencies of T cells in the female genital tract (n=126). Non-parametric Spearman correlations and Kruskal-Wallis statistical tests were performed. Statistical analyses were performed with false-discovery rate correction using the step-down procedure where appropriate. p<0.05 was considered statistically significant.

We next compared cytokines and T cell activation across exposure groups. While cytokine concentrations did not differ significantly between groups (data not shown), AGYW with evidence of past, resolved infection had significantly lower frequencies of activated cervical CD4+CD38+ T cells (median: 56.6%) compared to those with a recent, primary infection (p=0.018; median: 70%) or with untreated/recurrent infection (p=0.022; median: 77.2%; **Figure 2D**). Furthermore, women with untreated/recurrent exposure had significantly higher frequencies of activated cervical CD4+CD38+HLA-DR+ T cells compared to unexposed AGYW (medians: 32.7% and 17.75%, p=0.018) or those who had cleared infection (medians: 32.7% and 16.1%, p=0.022; **Figure 2E**). The observed association between cumulative *C. trachomatis* exposure and elevated cervical T cell activation underscores the potential for immune-mediated pathology and suggests sustained mucosal immune stimulation, possibly reflecting T cell recruitment to the genital tract in response to persistent antigenic challenge.

### Circulating *C. trachomatis*-Specific CD4+ T Cells reflect Level of Exposure

We measured systemic *C. trachomatis*-specific CD4+ T cell cytokine responses in peripheral blood mononuclear cells (PBMCs) from AGYW who were either NAAT positive or seropositive at baseline (n=46; **Figure 3A**). Most women (n=36/46) had detectable CD4+ T cell responses to rMOMP, regardless of NAAT status (**Figure 3B**). We also found no statistical differences in the number of NAAT-or NAAT+ participants producing any cytokine (75% and 79% respectively; p>0.99), IFN-γ (63% and 58%, respectively; p>0.99), IL-2 (63% and 47%, respectively p=0.699) or TNF (50% for both; p>0.99; **Figure 3B**). Similarly, there were no statistical differences between T cell responses in women who were NAAT- and NAAT+ in terms of CD4+ T cells producing any cytokine (median responses: 0.13% and 0.06%, respectively; p=0.378), or producing IFN-γ (median responses: 0.069% and 0.034%, respectively; p=0.383), IL-2 (median responses: 0.082% and 0.06%, respectively; p=0.321) or TNF (median of responses: 0.086% and 0.06%, respectively; p=0.771; **Figure 3B**). However, when participants were stratified by exposure, clear differences in responses emerged. All AGYW with untreated or recurrent infection had detectable CD4+ T cell responses (100%), compared to 67% of those with a primary exposure (p=0.017), although the last group was very small (n=8). Nevertheless, the magnitude of the T cell response was significantly lower in those with or recurrent infection (median response: 0.05%) compared to those with primary (median response: 0.12%; p=0.01) or cleared infection (median response: 0.13%; p=0.01; Figure 3C). There were no significant differences in cytokine-specific responses (p>0.05; IFN-γ, IL-2, or TNF) across exposure groups (**Figure 3D**).

**Figure 3.**
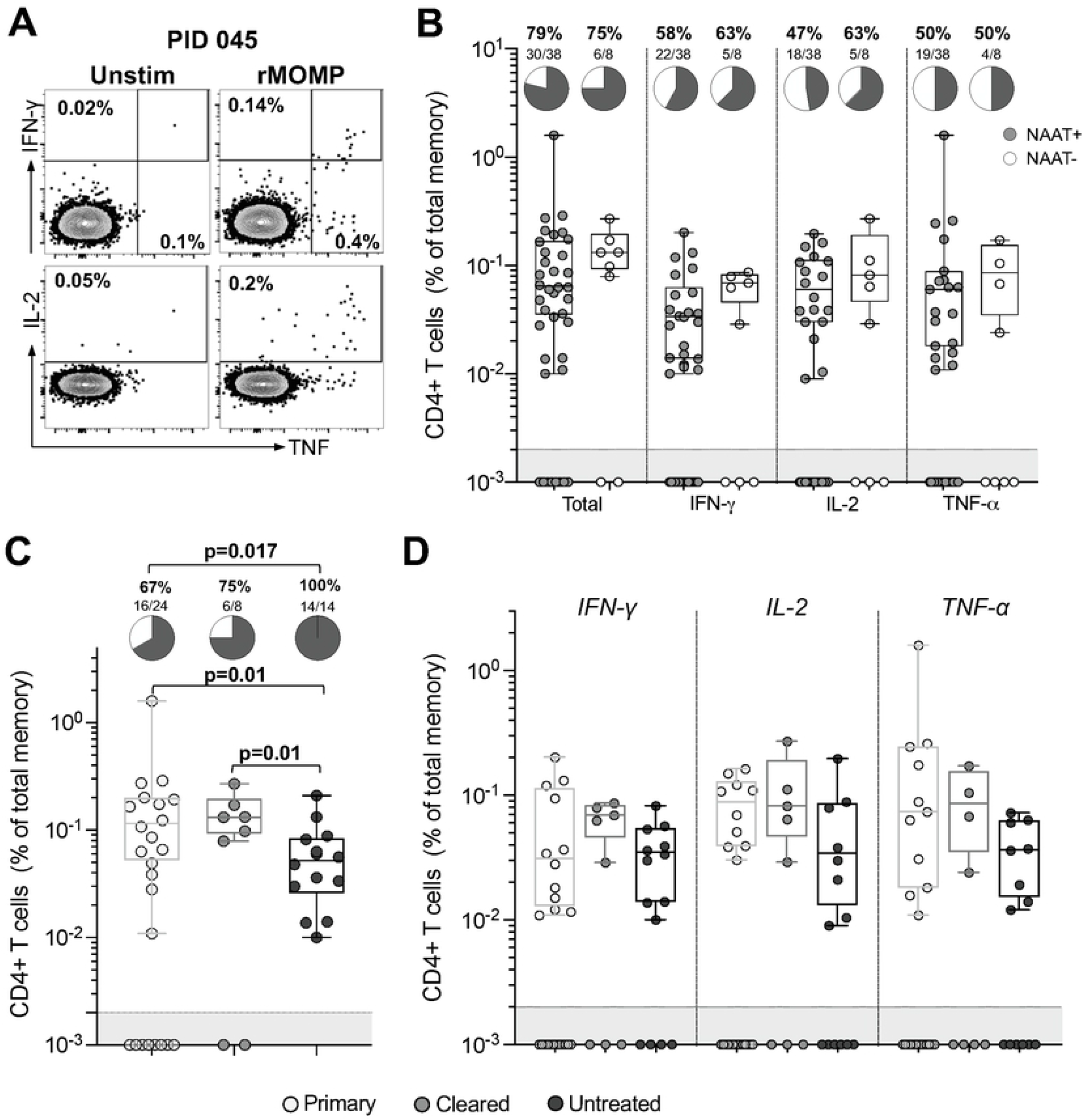
Peripheral CD4+ T cell responses to *C. trachomatis* are detectable in highly exposed AGYW. (A) Representative flow cytometry plots showing the IFN-γ, IL-2 and TNF response to rMOMP in one individual. (B) Frequency of CD4^+^ T cells responding to rMOMP in participants with a positive or negative NAAT result at baseline (n=46). The number of participants with a positive cytokine response is represented by the pie charts. (C) Frequency of CD4^+^ T cells producing any cytokine in response to rMOMP across the three exposure groups (n=46). Pie charts indicate the number of participants with a positive cytokine response defined as twice the background value and a net response >0.005%. (D) *C. trachomatis-*specific CD4+ T cells producing IFN-γ, IL-2 or TNF. The box and whisker plots represent positive cytokine responses only. Each symbol represents an individual. Data were analysed using a Kruskal-Wallis test. Fisher’s exact test was used to determine differences between pie graphs. p<0.05 was considered statistically significant.

We next investigated the functional quality of the CD4+ T cell response to *C. trachomatis* by examining cytokine co-expression. Cytokine profiles were significantly different between AGYW with different exposures (p<0.0001; **Figure 4**). Those experiencing a primary infection had predominantly monofunctional CD4+ T cells (median: 67.5%, IQR: 51-98.8%), and a low frequency of CD4+ T cells producing three cytokines (median: 6.62%, IQR: 0-19.9%). In contrast, AGYW with cleared infection had a more diverse response, with roughly equal proportions of T cells producing one (medians and IQR: 40.5%; 15.9-86.3%) or two cytokines (40.2%; 9.1-45.4%), and a median of 18.2% (IQR: 4.6-35.4%) of cells producing all three measured cytokines. Interestingly, participants with untreated/recurrent infection lacked these highly functional T cells producing three cytokines (median: 0%, IQR: 0-5.9%), and showed a predominance of dual-cytokine-producing cells (median: 43.8%, IQR: 21-100%). These findings suggest that while persistent exposure to *C. trachomatis* increases the likelihood of detecting systemic T cell responses, it may also be associated with diminished functional quality, particularly a lack of cells co-expressing IFN-γ, IL-2 and TNF. This highlights a potential link between chronic antigenic stimulation and T cell exhaustion or skewing, with implications for protective immunity and vaccine design.

**Figure 4.**
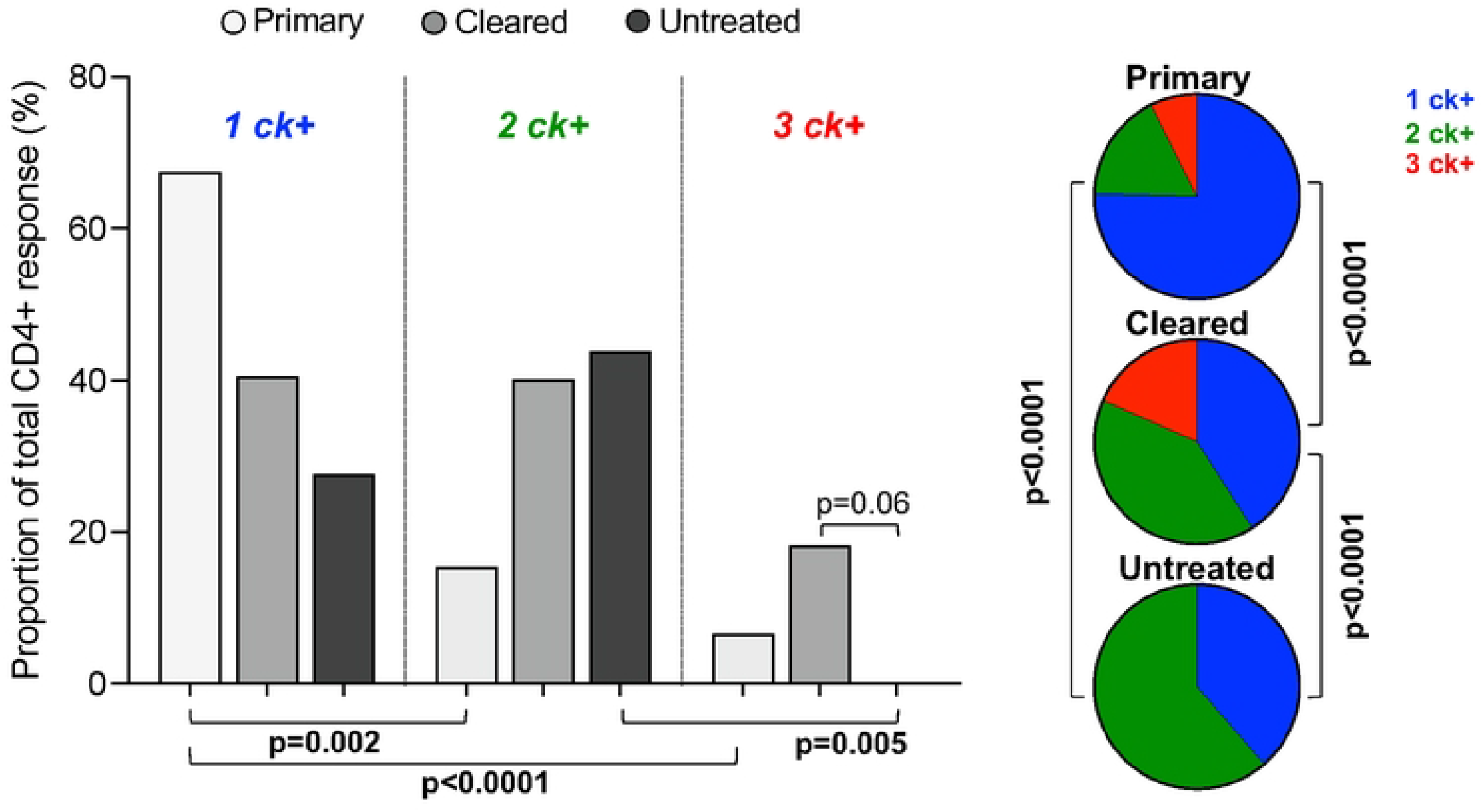
Exposure to *C. trachomatis* is reflected in CD4+ T cell function. The graph shows the proportion of *C. trachomatis-*specific CD4+ T cells producing different combinations of cytokines (n=36). The pies show the polyfunctionality profiles of *C. trachomatis-*specific CD4+ T cells stratified by exposure (n=36). Data shown are from participants with a positive total cytokine response. Each bar represents the median. Data were analysed using a Kruskal-Wallis test. Fisher’s exact test was used to determine differences between pie graphs. p<0.05 was considered statistically significant.

### *C. trachomatis*-specific T Cells are Associated with Reduced Genital Inflammation

To understand how *C. trachomatis*-specific T cell immunity might influence pathogen-associated immunopathology in the genital tract, we examined relationships between peripheral CD4+ T cell responses and cervicovaginal cytokine concentrations in AGYW with matched data. Notably, the frequency of CD4+ T cells producing IFN-γ in response to *C. trachomatis* stimulation was inversely correlated with levels of several proinflammatory mediators in the genital tract, including IL-1β (p=0.007, *r*=–0.397), TNF (p=0.028, *r*=–0.328), MIP-1β (p=0.045, *r*=–0.301), and IL-17 (p=0.024, *r*=– 0.337; **Figure 5**). Similarly, the frequency of *C. trachomatis*-specific CD4+ T cells producing TNF was significantly negatively associated with cervicovaginal IL-12p70 (p=0.039, *r*=–0.308). Importantly, none of these mucosal cytokines were associated with exposure, suggesting that these correlations reflect antigen-specific immune regulation rather than cumulative exposure alone.

**Figure 5.**
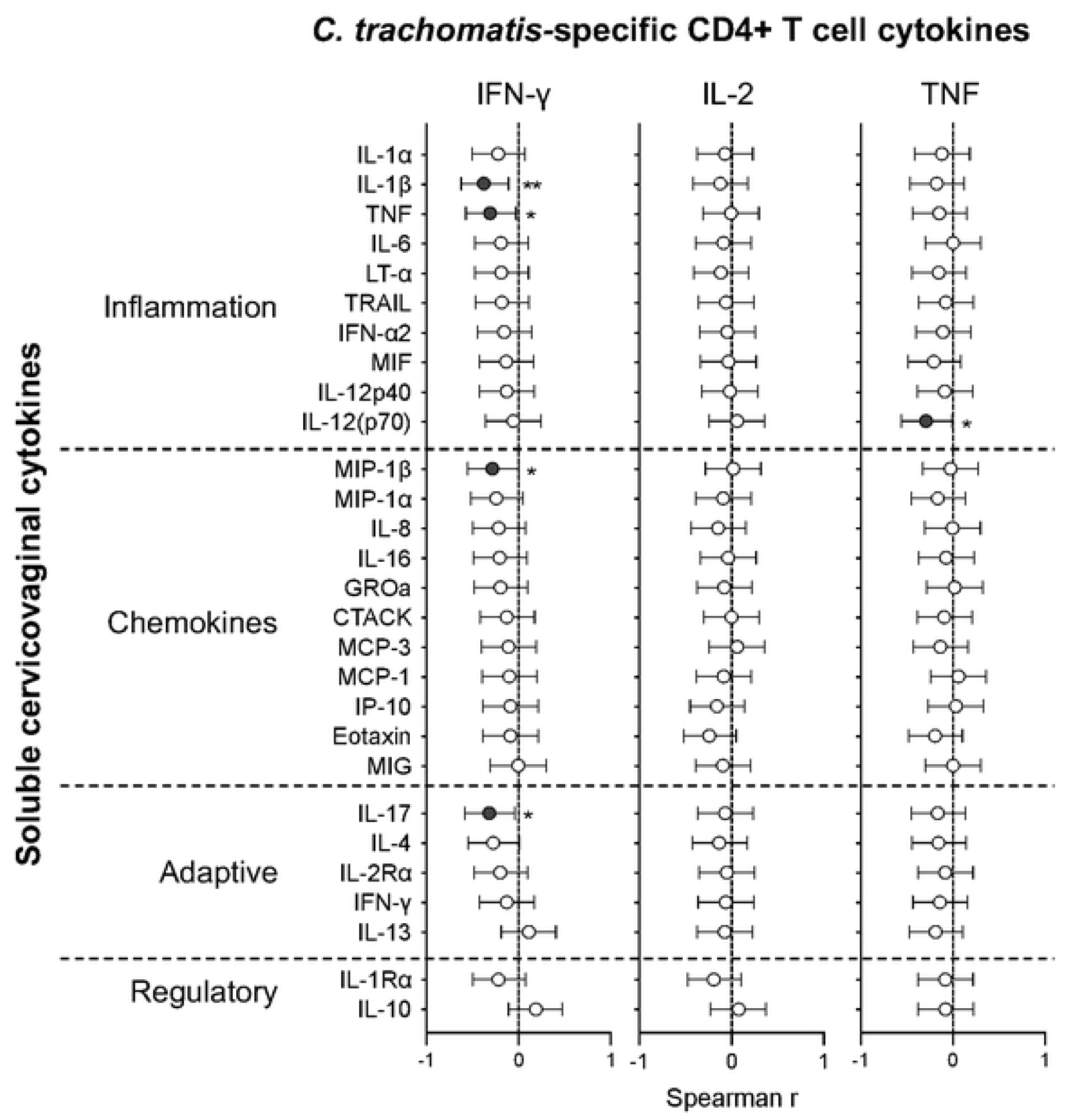
Associations between peripheral *C. trachomatis-specific* CD4+ T cell responses and the female genital tract immune milieu. Spearman correlation coefficient (r) and the 95% confidence interval are shown for associations between *C. trachomatis-*specific CD4+ T cells producing IFN-γ, IL-2 or TNF, and cervicovaginal cytokine concentrations (n=22). Filled circles indicate statistically significant correlations. p<0.05 was considered statistically significant. ^*^p<0.05; ^**^ p<0.01; ^***^ p<0.001.

## DISCUSSION

*Chlamydia trachomatis* is associated with poor reproductive health outcomes in women, contributing to infertility, pelvic inflammatory disease, and HIV acquisition risk [15–17,33]. Infections are often asymptomatic and undiagnosed in developing countries, where syndromic management is standard-of-care. Chlamydia remains a major health concern, with vaccines urgently required. Although there is some evidence for partial protection after natural infection, this is poorly defined, highlighting the need to understand chlamydia immunity in the context of its natural history. We examined circulating *C. trachomatis*-specific CD4+ T cells and their relationship with the genital tract immune milieu in women who had primary infection, untreated/recurrent infection, or cleared/treated prior infection.

Several important observations emerged from women with high *C. trachomatis* exposure in our study. Untreated/recurrent infections (NAAT+IgG+) were associated with activated cervical T cells, consistent with previous studies describing an increased risk of inflammatory sequelae after recurrent *C. trachomatis* infection [20,34,35]. In terms of *C. trachomatis*-specific T cell immunity, peripheral blood responses were detected in all women with untreated/recurrent infections, likely due to prolonged or repeated antigen exposure. Notably, we also found evidence of immune defects in the CD4+ T cell response to *C. trachomatis* in these women. These highly exposed women had significantly lower frequencies of *C. trachomatis*-specific Th1 responses compared to women with a primary or previous infection. Furthermore, there was a near-total absence of highly functional *C. trachomatis*-specific CD4+ T cells simultaneously producing three cytokines in women with recurrent infection. Meanwhile, women with cleared infections had the highest proportion of these cells. These important and novel findings emphasise that both the magnitude and quality of the CD4+ T cell response are vital for protective immunity to *C. trachomatis*.

These findings may also reflect underlying T cell trafficking dynamics. In women with long-term *C. trachomatis* exposure, antigen-specific CD4+ T cells may have migrated from peripheral blood to the site of infection, contributing to local immunity. Supporting this, murine studies have shown that *C. trachomatis*-specific T cells producing IFN-γ circulate between the genital tract and peripheral blood [36]. Clinical studies have similarly reported elevated IFN-γ concentrations in the female genital tract during recurrent *C. trachomatis* infections [32,37]. Here, cervicovaginal IFN-γ concentrations correlated with anti-chlamydia IgG titres, reinforcing its role as a marker of cumulative exposure. Given the well-established role of IFN-γ in pathogen clearance, we hypothesise that the cells producing IFN-γ, including *C. trachomatis*-specific Th1 cells, contribute to protective immunity at mucosal surfaces.

Consistently, we found significant negative associations between the frequency of peripheral *C. trachomatis*-specific CD4+ T cells and concentrations of cervicovaginal inflammatory mediators like IL-1β and TNF, indicative of a reduction in potential immunopathology when higher circulating antigen-specific T cells are detected. These findings suggest that systemic *C. trachomatis*-specific Th1 responses may not only reflect effective immune control but also actively contribute to the regulation of mucosal inflammation, potentially limiting immunopathology. Importantly, these associations were independent of exposure, highlighting the contribution of *C. trachomatis*-specific Th1 responses in shaping mucosal immune homeostasis, regardless of exposure history. Interestingly, women with untreated or recurrent infections had few three-function CD4+ T cells, despite a high prevalence of detectable responses. This lack of functional diversity may reflect T cell exhaustion or impaired differentiation, processes commonly associated with chronic antigenic stimulation in other infectious settings [38].

We classified participants into exposure groups using baseline NAAT and serology to gain a more nuanced understanding of *C. trachomatis* infection history. However, there were some caveats to this approach. Since only 17/122 AGYW reported a previous, unspecified STI, we surmised that most NAAT+ seropositive participants were untreated for *C. trachomatis*, rather than experiencing persistent infection due to treatment failure. Given the lack of treatment data, we cannot rule out persistent infections in all individuals. Nevertheless, these groupings still reflect the relative level of *C. trachomatis* exposure and provide valuable insights into chronic antigen exposure and immunity. A limitation of this study was that NAAT was only performed on vulvovaginal swabs, potentially missing anorectal infections [39] which would affect the classification of NAAT-participants.

T cell responses were assessed using a single antigen, the major outer membrane protein (MOMP), which is highly immunogenic [40], but may not fully capture the breadth of the T cell response. Recent research shows that T cell responses to other proteins e.g. heat-shock protein 60, may also play a role in host immunity [25,41,42]. T cell responses to C. trachomatis occur at low frequencies, making them difficult to detect in humans [25,26,42]. Although technically challenging, C. trachomatis-specific T cells should ideally be studied at the site of infection to fully disentangle the development of protective immunity and immunopathology. Given the focus on Th1 responses, particularly IFN-γ, due to its key role in immunity to C. trachomatis, mucosal-acting cytokines like IL-17 and IL-22 or immunoregulatory cytokines such as IL-10, may go unappreciated. In future, identifying a robustly detectable response-ideally in blood-will be crucial for evaluating vaccine-induced immunity and identifying potential correlates of protection.

In conclusion, we showed that women with untreated or recurrent infections had lower frequencies of *C. trachomatis*-specific CD4+ T cells, and a smaller proportion of these cells were highly functional. Furthermore, peripheral T cell responses negatively correlated with inflammation in the genital tract. Our data clearly indicate that both the magnitude and function of the CD4+ T cell immune response to *C. trachomatis* are important components of protective immunity. These findings underscore the importance of including mucosal immune markers and functional systemic responses in the evaluation of candidate chlamydia vaccines, especially in high-risk populations such as AGYW.

## MATERIALS AND METHODS

### Study participants

AGYW (16-22 years old) from South Africa were enrolled in the Women’s Initiative in Sexual Health (WISH) study and provided samples every two months for six months [29,31]. A total of n=149 AGYW completed the baseline study visit, 127/149 completed two visits and 88/149 completed three. Participants were excluded if they were living with HIV, pregnant, menstruating at time of sampling, or if they were sexually active, douched or used spermicides in the last two days, or took antibiotics in the last two weeks. All participants gave written, informed assent/consent to be involved in the study. For participants under the age of 18, written consent was obtained from the legal guardian. The Human Research Ethics Committee of the University of Cape Town (267/2013) approved the study.

### Study procedures

Samples were collected at each study visit [29,32]. A vulvovaginal swab was collected for STI testing, cervicovaginal secretions (CVS) were collected through insertion of a menstrual cup (Softcup; The Flex Company, USA) into the vagina for 30 minutes. Participants were tested for STIs at every visit. Specimens positive for *C. trachomatis* were also tested for lymphogranuloma venereum (LGV)-associated L1-L3 genovars. All participants were negative for LGV. Participants who had signs/symptoms of STIs, or tested positive, were offered treatment and a partner referral letter.

### Sample processing

Peripheral blood mononuclear cells (PBMC) and plasma were isolated by density gradient centrifugation with Histopaque (Sigma Aldrich, USA) within 4 hours of collection. Cells were cryopreserved in foetal calf serum (HyClone; Cytiva, USA) with 10% DMSO (Sigma-Adrich, USA) and stored in liquid nitrogen until use. Plasma was aliquoted and stored at -80^°^ C. Cervicovaginal secretions were removed from the menstrual cup as previously described [29,43], diluted 1:5 with phosphate buffered saline (PBS; Sigma Alrich, USA), aliquoted, and stored at -80^°^ C.

### Stimulation of T cells

We selected participants with a positive *C. trachomatis* nucleic acid amplification test (NAAT) or a who were seropositive for *C. trachomatis* at any visit during the six-month study period, where samples were available (n=46). Cryopreserved PBMC from the baseline visit were thawed and rested in R10 medium (Gibco RPMI 1640; Thermo Fisher Scientific, USA), with the addition of 10% heat-inactivated FCS (HyClone; Cytiva, USA) and 50U/mL of penicillin-streptomycin (Sigma-Aldrich, USA) for 4 hours prior to stimulation. Cells were incubated in R10 with rMOMP protein (10μg/mL; Uniprot Accession #P17451.1, ProMab, USA) for 18 hours at 37°C in the presence of the co-stimulatory antibodies anti-CD28 and anti-CD49d (1μg/mL each; BD Biosciences, USA). Brefeldin A (5μg/mL, Sigma-Aldrich, USA) was added after 3 hours. Phorbol 12-myristate 13-acetate (0.04μg/mL, Sigma-Aldrich, USA) and ionomycin (100μg/mL, Sigma-Aldrich, USA) were used as a positive control. Unstimulated cells were included as a negative control.

### Multiparameter flow cytometry

Stimulated PBMC were stained with a violet viability dye (ViViD; Molecular Probes, USA), followed by staining with CD14-Pacific blue, CD19-Pacific blue, CD4-PE-Cy5.5 (all Invitrogen, USA), CD8-BV711 (Biolegend, USA), CD27-PE-Cy-5 (Beckman Coulter, USA) and CD45RO-BV785 (Biolegend, USA). Cells were permeabilised with Cytofix-Cytoperm (BD Biosciences, USA), stained intracellularly with CD3-APC-H7, IFN-γ-Alexa700, IL-2-APC (all from BD Biosciences, USA) and TNF-α-PE-Cy7 (Biolegend, USA), and fixed with CellFix (BD Biosciences, USA). Cells were acquired on a BD LSRII (BD Biosciences, USA), using FACSDiva software. Data were analysed using FlowJo v10 (TreeStar, BD Biosciences, USA) and Pestle and Spice [44]. The gating strategy is shown in **Supplementary Figure 1**. A positive cytokine response was defined as at least double the background frequency of the unstimulated sample, and a net response >0.005%. All data are reported after background subtraction.

### Measurement of *C. trachomatis*-specific IgG by ELISA

Baseline visit plasma from 145 women were used to measure IgG using a semi-quantitative ELISA according to manufacturer’s instructions (EuroImmun; Germany). Briefly, plasma was thawed overnight at 4° C, diluted and incubated on the supplied pre-coated plate. The plate was washed, and incubated with peroxidase labelled anti-IgG, followed by a final incubation with tetramethylbenzidine (TMB) substrate. Stop solution was added and colour intensity measured at 450nm and 650nm within 15 minutes. The OD reading at 650nm (reference wavelength) was subtracted from the reading at 450nm. The standard curve was used to calculate the relative units (RU) of IgG. Results of >22 RU/ml were positive, 16-22RU/ml were borderline, and <16 RU/ml was considered negative. Trachoma (serovars A-C) is not endemic to South Africa, and all participants were negative for LGV, suggesting the IgG detected arose from *C. trachomatis* serovar D-K infections. Similarly, cross-reactive antibodies generated by *C. pneumoniae* are unlikely due to their rarity in South Africa [45,46], and the species specificity of the kit.

### Cervicovaginal cytokine measurement

Diluted cervicovaginal secretions were thawed and filtered through a 0.2μm cellulose acetate filter. The Human Cytokine Group I 27-plex and Human Cytokine 21-plex kits (Bio-Rad, USA) were used to quantify a range of 44 cytokines, chemokines and growth factors. Values below the detection limit were recorded as half of the lowest measured concentration for each cytokine. IL-2, IL-5, IL-15 and RANTES were excluded for all analyses as the values did not pass QC or >50% of values were below the detection limit.

## Statistical Analysis

Statistical analyses were performed using Prism 9 (GraphPad, USA). Non-parametric tests (Mann-Whitney U, Wilcoxon matched pairs, Spearman Rank) were used for comparisons. Fischer’s exact test was used to compare proportions of responders. A p value of <0.05 was considered statistically significant.

## ACKNOWLEDGEMENTS

We thank the study participants for providing samples and for their time and commitment to the study; the WISH clinical study team particularly Ms Penelope Ngcobo and Sr Janine Nixon for their essential contribution; and Mrs Kathryn Norman, for administrative assistance.

## CONFLICT OF INTEREST

The authors have no conflict of interest.

## FUNDING

This work was supported by the European and Developing Countries Clinical Trials Partnership (EDCTP) 1 (SP-2011-41304-038 to J-A.S.P), and the Fondation Botnar under the EDCTP2 programme (TMA2020CDF-3187 to R.B.) supported by the European Union; the South African Medical Research Council (SAMRC; self-initiated research award to J-A.S.P.); the Carnegie Corporation of New York (postdoctoral fellowship to R.B.). The Desmond Tutu Health Foundation recognizes the support from ViiV health care in their Youth Shield program. S.D. received funding from the National Research Foundation (NRF) and Poliomyelitis Research Foundation (PRF). S.B. was funded by the HIV Vaccine Trials Network SHAPe Program, the Fogarty Foundation and the SAMRC. The funders had no role in study design, data collection and analysis, decision to publish, or preparation of the manuscript.

## AUTHOR CONTRIBUTIONS

H.B.J. and J-A.S.P. designed and led the WISH study. L-G.B., K.G. and S.B. recruited the cohort. V.M. and D.A.L. oversaw and performed the STI testing. H.G. provided technical expertise. R.B. and J-A.S.P. designed the substudy and experiments. R.B., M.L., S.D., S.B. and S.Z.J. performed the experiments. R.B., S.D. and S.B. analysed the data. R. B., M.L. and J-A.S.P. wrote the manuscript. All authors reviewed manuscript drafts and approved the final manuscript.

## Notes

### Competing Interest Statement

The authors have declared no competing interest.

## REFERENCES

1. WHO (2024) Chlamydia Fact Sheet. Available: https://www.who.int/news-room/fact-sheets/detail/chlamydia. xAccessed 5 March 2025.

2. Torrone EA, Morrison CS, Chen P-L, Kwok C, Francis SC, et al. (2018) Prevalence of sexually transmitted infections and bacterial vaginosis among women in sub-Saharan Africa: An individual participant data meta-analysis of 18 HIV prevention studies. PLOS Med 15: e1002608. Available: 10.1371/journal.pmed.1002608.

3. Pinto CN, Niles JK, Kaufman HW, Marlowe EM, Alagia DP, et al. (2021) Impact of the COVID-19 Pandemic on Chlamydia and Gonorrhea Screening in the U.S. Am J Prev Med 61: 386–393. Available: 10.1016/j.amepre.2021.03.009.

4. Du M, Yan W, Jing W, Qin C, Liu Q, et al. (2022) Increasing incidence rates of sexually transmitted infections from 2010 to 2019: an analysis of temporal trends by geographical regions and age groups from the 2019 Global Burden of Disease Study. BMC Infect Dis 22: 1–16. Available: 10.1186/s12879-022-07544-7.

5. Zheng Y, Yu Q, Lin Y, Zhou Y, Lan L, et al. (2022) Global burden and trends of sexually transmitted infections from 1990 to 2019: an observational trend study. Lancet Infect Dis 22: 541–551. doi:10.1016/S1473-3099(21)00448-5.

6. Detels R, Green AM, Klausner JD, Katzenstein D, Gaydos C, et al. (2011) The incidence and correlates of symptomatic and asymptomatic chlamydia trachomatis and neisseria gonorrhoeae infections in selected populations in five countries. Sex Transm Dis 38: 503–509. doi:10.1097/OLQ.0b013e318206c288.

7. Kularatne RS, Niit R, Rowley J, Kufa-Chakezha T, Peters RPH, et al. (2018) Adult gonorrhea, chlamydia and syphilis prevalence, incidence, treatment and syndromic case reporting in South Africa: Estimates using the Spectrum-STI model, 1990-2017. PLoS One 13: 1–22. doi:10.1371/journal.pone.0205863.

8. Dhouib W, Zemni I, Kacem M, Bennasrallah C, Fredj M Ben, et al. (2020) Syndromic Management of Female Sexually Transmitted Infections at Primary Care Level. A Descriptive Cross-Sectional Study Tunisia. Res Sq: 1–13. doi:10.21203/rs.3.rs-71576/v1.

9. Kaida A, Dietrich JJ, Laher F, Beksinska M, Jaggernath M, et al. (2018) A high burden of asymptomatic genital tract infections undermines the syndromic management approach among adolescents and young adults in South Africa: Implications for HIV prevention efforts. BMC Infect Dis 18: 1–11. doi:10.1186/s12879-018-3380-6.

10. Brunham RC, Rappuoli R (2013) Chlamydia trachomatis control requires a vaccine. Vaccine 31: 1892–1897. Available: 10.1016/j.vaccine.2013.01.024.

11. Abraham S, Juel HB, Bang P, Cheeseman HM, Dohn RB, et al. (2019) Safety and immunogenicity of the chlamydia vaccine candidate CTH522 adjuvanted with CAF01 liposomes or aluminium hydroxide: a first-in-human, randomised, double-blind, placebo-controlled, phase 1 trial. Lancet Infect Dis 3099: 1–10. Available: https://linkinghub.elsevier.com/retrieve/pii/S1473309919302798.

12. Zhong G, Brunham RC, de la Maza LM, Darville TL, Deal C (2017) National institute of allergy and infectious diseases workshop report: “Chlamydia vaccines: The way forward”. Vaccine. Available: http://linkinghub.elsevier.com/retrieve/pii/S0264410X17314834.

13. Brunham RC (2021) Problems With Understanding Chlamydia trachomatis Immunology. J Infect Dis: 1–16. doi:10.1093/infdis/jiab610.

14. Qu Y, Frazer LC, O’Connell CM, Tarantal AF, Andrews CW, et al. (2015) Comparable Genital Tract Infection, Pathology, and Immunity in Rhesus Macaques Inoculated with Wild-Type or Plasmid-Deficient Chlamydia trachomatis Serovar D. Infect Immun 83: 4056–4067. doi:10.1128/IAI.00841-15.

15. Beatty WL, Byrne GI, Morrison RP (1994) Repeated and persistent infection with Chlamydia and the development of chronic inflammation and disease. Trends Microbiol 2: 94–98. Available: http://www.ncbi.nlm.nih.gov/pubmed/8156277.

16. Darville T, Hiltke TJ (2010) Pathogenesis of Genital Tract Disease Due to Chlamydia trachomatis. J Infect Dis 201: 114–125. doi:10.1086/652397.

17. Masson L, Passmore JAS, Liebenberg LJ, Werner L, Baxter C, et al. (2015) Genital Inflammation and the Risk of HIV Acquisition in Women. Clin Infect Dis 61: 260–269. doi:10.1093/cid/civ298.

18. Arno JN, Katz BP, McBride R, Carty GA, Batteiger BE, et al. (1994) Age and clinical immunity to infections with Chlamydia trachomatis. Sex Transm Dis 21: 47–52. Available: 10.1371/journal.pone.0192357.

19. Batteiger BE, Tu W, Ofner S, Van Der Pol B, Stothard DR, et al. (2010) Repeated Chlamydia trachomatis Genital Infections in Adolescent Women. J Infect Dis 201: 42–51. doi:10.1086/648734.

20. Bautista CT, Hollingsworth BP, Sanchez JL (2018) Repeat Chlamydia Diagnoses Increase the Hazard of Pelvic Inflammatory Disease among US Army Women: A Retrospective Cohort Analysis. Sex Transm Dis 45: 770–773. doi:10.1097/OLQ.0000000000000878.

21. Wijers JNAP, Dukers-Muijrers NHTM, Hoebe CJPA, Wolffs PFG, Van Liere GAFS (2020) The characteristics of patients frequently tested and repeatedly infected with Chlamydia trachomatis in Southwest Limburg, the Netherlands. BMC Public Health 20: 1–8. doi:10.1186/s12889-020-09334-9.

22. Lampe MF, Wilson CB, Bevan MJ, Starnbach MN (1998) Gamma interferon production by cytotoxic T lymphocytes is required for resolution of Chlamydia trachomatis infection. Infect Immun 66: 5457–5461. Available: http://www.ncbi.nlm.nih.gov/entrez/query.fcgi?cmd=Retrieve&db=PubMed&dopt=Citation&list_uids=9784557.

23. Gondek DC, Olive AJ, Stary G, Starnbach MN (2012) CD4+ T Cells Are Necessary and Sufficient To Confer Protection against Chlamydia trachomatis Infection in the Murine Upper Genital Tract. J Immunol 189: 2441–2449. Available: http://www.ncbi.nlm.nih.gov/pubmed/17977024.

24. Helble JD, Starnbach MN (2021) T cell responses to Chlamydia. FEMS Microbiol Rev.

25. Russell AN, Zheng X, O’Connell CM, Wiesenfeld HC, Hillier SL, et al. (2016) Identification of Chlamydia trachomatis Antigens Recognized by T Cells From Highly Exposed Women Who Limit or Resist Genital Tract Infection. J Infect Dis 214: 1884–1892. Available: http://jid.oxfordjournals.org/content/early/2016/10/12/infdis.jiw485.full.pdf.

26. Jordan SJ, Gupta K, Ogendi BMO, Bakshi RK, Kapil R 4, et al. (2017) The Predominant CD4 + Th1 Cytokine Elicited to Chlamydia trachomatis Infection in Women is Tumor Necrosis Factor-Alpha, not Interferon-Gamma. Vaccine Immunol 1. doi:10.1128/CVI.00010-17.

27. Jordan SJ, Bakshi RK, Brown LDT, Chi X, Geisler WM (2018) Stimulated peripheral blood mononuclear cells from chlamydia-infected women release predominantly Th1-polarizing cytokines. Cytokine: 0–1. Available: 10.1016/j.cyto.2018.06.017.

28. Cohen CR, Koochesfahani KM, Meier AS, Shen C, Karunakaran K, et al. (2005) Immunoepidemiologic profile of Chlamydia trachomatis infection: importance of heat-shock protein 60 and interferon-gamma. J Infect Dis 192: 591–599. Available: http://www.ncbi.nlm.nih.gov/pubmed/16028127%5Cnhttp://jid.oxfordjournals.org/content/192/4/591.full.pdf.

29. Dabee S, Barnabas SL, Lennard KS, Jaumdally SZ, Gamieldien H, et al. (2019) Defining characteristics of genital health in South African adolescent girls and young women at high risk for HIV infection. PLoS One 14: e0213975. Available: 10.1371/journal.pone.0213975.

30. Lennard K, Dabee S, Barnabas SL, Havyarimana E, Blakney A, et al. (2017) Microbial composition predicts genital tract inflammation and persistent bacterial vaginosis in adolescent South African women. Infect Immun: IAI.00410-17. Available: http://iai.asm.org/lookup/doi/10.1128/IAI.00410-17.

31. Barnabas SL, Dabee S, Passmore J-AS, Jaspan HB, Lewis DA, et al. (2017) Converging epidemics of sexually transmitted infections and bacterial vaginosis in southern African female adolescents at risk of HIV. Int J STD AIDS: 095646241774048. Available: http://journals.sagepub.com/doi/10.1177/0956462417740487.

32. Dabee S, Barnabas S, Kullin B, Versteeg B, Mkhize N, et al. (2025) Cervicovaginal inflammation and HIV target cell activation in adolescent girls and young women with asymptomatic. medRxiv Prepr. doi:10.1101/2025.02.04.25321658.

33. Darville T (2021) Pelvic Inflammatory Disease Due to Neisseria gonorrhoeae and Chlamydia trachomatis: Immune Evasion Mechanisms and Pathogenic Disease Pathways. J Infect Dis 224: S39– S46. doi:10.1093/infdis/jiab031.

34. den Heijer CDJ, Hoebe CJPA, Driessen JHM, Wolffs P, van den Broek IVF, et al. (2019) Chlamydia trachomatis and the Risk of Pelvic Inflammatory Disease, Ectopic Pregnancy, and Female Infertility: A Retrospective Cohort Study Among Primary Care Patients. Clin Infect Dis: 1–8. Available: https://academic.oup.com/cid/advance-article/doi/10.1093/cid/ciz429/5554161.

35. Reekie J, Donovan B, Guy R, Hocking JS, Kaldor JM, et al. (2018) Risk of Pelvic Inflammatory Disease in Relation to Chlamydia and Gonorrhea Testing, Repeat Testing, and Positivity: A Population-Based Cohort Study. Clin Infect Dis 66: 437–443. doi:10.1093/cid/cix769.

36. Labuda JC, Pham OH, Depew CE, Fong KD, Lee B, et al. (2021) Circulating immunity protects the female reproductive tract from Chlamydia infection. Proc Natl Acad Sci U S A 118. doi:10.1073/pnas.2104407118.

37. Agrawal T, Vats V, Wallace PK, Salhan S, Mittal A (2007) Cervical cytokine responses in women with primary or recurrent chlamydial infection. J Interf Cytokine Res 27: 221–226. Available: http://www.liebertonline.com/doi/abs/10.1089/jir.2006.0132.

38. Wherry EJ, Kurachi M (2015) Molecular and cellular insights into T cell exhaustion. Nat Rev Immunol 15: 486–499. doi:10.1038/nri3862.

39. Peters RPH, Dubbink JH, Van Der Eem L, Verweij SP, Bos MLA, et al. (2014) Cross-sectional study of genital, rectal, and pharyngeal chlamydia and gonorrhea in women in rural South Africa. Sex Transm Dis 41: 564–569. doi:10.1097/OLQ.0000000000000175.

40. Olsen AW, Rosenkrands I, Holland MJ, Andersen P, Follmann F (2021) A Chlamydia trachomatis VD1-MOMP vaccine elicits cross-neutralizing and protective antibodies against C/C-related complex serovars. npj Vaccines 6: 1–11. Available: 10.1038/s41541-021-00312-9.

41. Cheong HC, Lee CYQ, Cheok YY, Shankar EM, Sabet NS, et al. (2018) tCPAF, HSP60 and MOMP antigens elicit pro-inflammatory cytokines production in the peripheral blood mononuclear cells from genital Chlamydia trachomatis-infected patients. Immunobiology. doi:10.1016/j.imbio.2018.10.010.

42. Li Y, Warren JA, Poston TB, Clutton G, Shaw FR, et al. (2024) Chlamydia trachomatis induces low-frequency, sustained CD4 T cell responses in most women, predominantly targeting chlamydial protease-like activity factor, CPAF. J Infect Dis. doi:10.1093/infdis/jiae443.

43. Jaumdally SZ, Masson L, Jones HE, Dabee S, Hoover DR, et al. (2018) Lower genital tract cytokine profiles in South African women living with HIV: influence of mucosal sampling. Sci Rep 8: 1–12. doi:10.1038/s41598-018-30663-8.

44. Roederer M, Nozzi JL, Nason MC (2011) SPICE: Exploration and analysis of post-cytometric complex multivariate datasets. Cytom Part A 79 A: 167–174. doi:10.1002/cyto.a.21015.

45. Moore DP, Baillie VL, Mudau A, Wadula J, Adams T, et al. (2021) The Etiology of Pneumonia in HIV-uninfected South African Children: Findings from the Pneumonia Etiology Research for Child Health (PERCH) Study. Pediatr Infect Dis J 40: S59–S68. doi:10.1097/INF.0000000000002650.

46. National Institute for Communicable Diseases (2017) Communicable Diseases. doi:10.5005/jp/books/14242_31.

